# Identification and Targeting Putative G-Quadruplex Sequences in *Candida glabrata*: A Route to Virulence and Pathogenesis Control?

**DOI:** 10.1101/2021.08.06.455387

**Authors:** Ankush Yadav

## Abstract

An increase in the number of *Candida* species that are resistant to antifungal medication and increases worldwide. Even individuals that are never exposed to antibiotics showing resistance to antifungal drugs. The increase in resistant candida species strains requires a search for novel targets for new antifungal agents. Preventing infection caused by candida species is a tremendous challenge in medicine. Although availability and use of antifungal drugs, disseminated candidiasis leads to a high mortality rate of about 40-60%, poor diagnosis, and improper disease management. Interest in G-quadruplexes as a therapeutic target has been increased in recent years, following the implication of this non-canonical G-quadruplex secondary structure in pathological diseases. However, G-quadruplex has been reported in many pathogens contributing to virulence and pathogenesis, including bacterial pathogens such as Staphylcoccus aureus and Enterococcus spp. Etc, viruses such as SARS-CoV2, HIV, HPV etc., and Fungi such as Candida species, Aspergillus fumigatus, etc. The present aim of the study is to identify and targeting G-Quadruplex forming sequences present in the *Candida glabrata*. PQSFinder (R Package) identified more than 5000 putative G-quadruplex forming sequences. Out of these, we have used PQS present in the SDH1 gene of Candida glabrata. It may be a key target to ameliorate the *C.glabrata* infection because it encodes a protein that plays a vital role in energy production in *C.glabrata* cells shown in the figure. The structure was built based on the already current structure that is telomeric G4 AGGG (TTAGGG)_3_ {PDB ID-4G0F} for identifying its stabilization by Gold carbine derivatives. Molecular docking and ADMET analysis show that Compound A has the highest binding affinity and has the best ADMET properties among the two compounds. The present study represents PQS in the SDH1 gene could be a novel antifungal target.

## 1. Introduction

Modern medicine has prevented millions of people from dying of common infections, cancers, and immunodeficiency viruses. However, patients are left vulnerable to secondary infections such as fungal infections when those lifesaving treatments are based on suppressing the human immune system [Pappas et al., 2018]. A healthy human body and the immune system are fine-tuned to keep yeast and fungi at bay. Still, in an immunocompromised state, several species in the *Candida* genus are quick to take advantage of the situation and spread throughout the human body and blood. Blood stage infections of *Candida* species can have a mortality rate of almost 70%, and many isolates of multidrug-resistant fungal strains are currently emerging [Fisher et al., 2018]. Isolates of the *Candida* species, first isolated in 1844, are already drug-resistant to several of the few antifungal agents available [Vautier et al., 2012]. Because fungal infections historically have not been as devastating or frequent as bacterial or protozoan infections, the development of antifungal drugs has not been prioritized [Fisher et al., 2018]. Fungi share many fundamental cellular processes antifungal drugs target the fungal cell wall, a defining feature of fungal cells [Gow, Large and Munro, 2017].

*Candida* species are generally harmless eukaryotic commensal yeast that belongs to the Ascomycota phylum and recovered from human, mammalian, and environmental sources. *Candida* species commonly reside in mammals as part of the normal microbial flora on the mucosal surfaces of the genitourinary and gastrointestinal tracts in the healthy person [Kumamoto, 2011]. They often cause systematic and mucosal infection only when the host’s immunity becomes compromised [Vincent et al., 2009].

When the balance between the yeast and the host is disturbed, or the host’s immune system is impaired. This *Candida* species typically become opportunistic or pathogenic. When the use of antibiotics wipes out bacterial populations, it frees up new niches for *Candida*. [Turner & Butler, 2014; Pappas et al., 2018, Dadar et al., 2018]. Candida species are classified as superficial infections when infecting the skin, respiratory system, oral cavity, and gastrointestinal tract. In contrast, invasive infection is characterized by severe conditions like candidemia, endocarditis, and meningitis [De Rosa et al., 2009].

For causing infection, candida must evade host defenses and multiply in the host. The *Candida* species must propagate to other tissues and organs in systematic infection. Gastrointestinal and skin barrier disruption can lead to deep organ candidiasis. *Candida albicans* are responsible for about 50% of candidiasis and, non-albicans *Candida* species are responsible for the rest of the Candida infection.

Nowadays, these non-albicans *Candida* species such as *Candida glabrata* are of great concern and emerging as opportunistic pathogens. *Candida glabrata* has been classified as a non-pathogenic saprophyte of normal microbial flora present in healthy individuals and rarely causes fungal infections in humans [Haley, 1961]. *C. glabrata* is the second most organism that cause candidiasis after *Candida albicans. C. glabrata* do not showing dimorphic characteristics. Instead, it is found as blastoconidia in commensal microbial flora or during pathogenesis. *C. glabrata* was identified as *Cryptococcus glabratus*, and this name was first changed to *Torulopsis glabrata* in 1894 and then again renamed *C. glabrata*. The present taxonomy for *C. glabrata* would be Kingdom-Fungi; Subkingdom-*Dikarya*; Phylum-*Ascomycota*; Subphylum-*Saccharomycotina*; Class-Saccharomycetes; Order-*Saccharomycetales*; Family-*Saccharomycetaceae*; Genus-*Nakaseomyces*; Clade-*Candida*; Species-*glabrata*. The genome of the *Candida glabrata* consists of 13 Chromosomes starting from A to M and about 12.3 megabase pairs in Length [Dujon et al., 2004]. It is assumed that *Candida glabrata* contains 5293 ORFs, out of which only 4.5% ORFs that is 238, are verified by experiments that code for the proteins. The Length of the chromosomes also varies from 4, 91,328 to 14, 55,689 bp, and chromosome A is the smallest in size, whereas chromosome M is the highest.

Preventing infection caused by candida species is a tremendous challenge in medicine. Although availability and use of antifungal drugs, disseminated candidiasis leads to a high mortality rate of about 40-60%, poor diagnosis, and improper disease management.

An increase in the number of *Candida* species that are resistant to antifungal medication and increases worldwide. Even individuals that are never exposed to antibiotics showing resistance to antifungal drugs. *Candida* species strains resistant to antifungals (such as echinocandins and fluconazole) are increasingly being identified. Their presence usually correlates with high echinocandin and azoles background usage in hospitals or intensive care units [Arendrup & Perlin, 2014, Pfaller et al., 2011]. *Candida* species can form a drug-resistant biofilm that is a vital factor for pathogenesis in humans. The majority of sessile cells [Ramage et al., 2012] and biofilms [Kuhn & Ghannoum, 2004] within candida species are less susceptible to antifungal medications than the planktonic cells. The increase in resistant candida species strains requires a search for novel targets for new antifungal agents.

G-quadruplexes forming sequences are conserved (evolutionary) drug targets. They are found in the essential regions of the genome of prokaryotes, eukaryotes, and viruses (Murat & Balasubramanian, 2014). G-quadruplexes are secondary structures formed in nucleic acid by the association of Guanines in G-rich sequences to form G-tetrad structures. G-quadruplex is enriched in the regulatory regions, especially the telomeric and promoter regions of prokaryotic and eukaryotic genomes (Varshney et al., 2020). Studies have indicated that G-quadruplexes also formed on the DNAs and RNAs template strands (Endoh et al., 2013). Interest in G-quadruplexes as a therapeutic target has been increased in recent years, following the implication of this non-canonical G-quadruplex secondary structure in pathological diseases such as cancer (Balasubramanian et al., 2011). However, now G-quadruplex has been reported in the vast number of pathogens contributing to virulence and pathogenesis including bacterial pathogens such as Staphylcoccus aureus, Enterocuccus spp. Etc, viruses such as SARS-CoV2, HIV, HPV etc., and Fungi such as Candida species, Aspergillus fumigatus, Cryptococcus neoformans etc. (Warner et al., 2021).

## 2. Methods and Materials

### 2.1 Prediction of G-quadruplex forming sequences in Candida Species

Sequences of *Candida glabrata* were extracted from the GenBank database available at NCBI (National Center for Biotechnology Information). (www.ncbi.nlm.nih.gov). We have used Putative G-quadruplex Prediction Tool (http://bsbe.iiti.ac.in/bsbe/ipdb/pattern2.php) to identify the G-quadruplex forming sequences and re-evaluated the results with other online and R programming based PGQ mining tools such as PQSFinder and QGRS Mapper. The Putative G-quadruplex Predictiion Tool algorithm exploits the following criteria to determine the PGQ sequences in *Candida glabrata*.

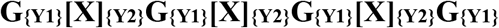

X= any nucleotide (A, G, C, T)
Y1= Length of consecutive guanine tract (Length vary between 2 to 7)
Y2= Length of Variable loop (loop length can range between 1 to 20)

All predicted PGQ sequences were recorded, and their location was elucidated using the Graphics mode of the GenBank database (http://www.ncbi.nlm.nih.gov/gene/).

### 2.2 Prediction of Secondary confirmation of SDH1 PQS and SDH1 G4 Structure

The hairpin conformation of SDH1 PQS was determined using an RNAfold web server. The entropy of each nucleotide that participated in the formation of hairpin conformation was elucidated by the RNAfold web server. The Structure of SDH1 G4 was not explained yet. Therefore the predicted model of SDH1 G4 was built based on the already present structure that is telomeric G4 AGGG (TTAGGG)_3_ {PDB ID-4G0F}. This sequence was selected because of its sequence similarity with the SDH1 G4 sequence. As represented in Table1, The 4G0F DNA sequence was modified to attain the nucleotide sequence of SDH1 G4 CGGG(CUCGGG)_3_ using the Discovery studio mutation tool. Adenine in position one was mutated to cytosine. The thymine residue in positions 5, 11, and 17 was mutated to cytosine, whereas thymine in positions 6,12, and 18 was mutated to uracil. Adenine in positions 7, 13, and 19 were also mutated to cytosine. The mutation was carried out in the loop residues, so that integrity of the G-tetrads was not compromised. The 2’ position of each ribose was modified from 2’-OH (oxy) to 2’-H (deoxy) in order to convert the nucleotide sequence into an RNA nucleotide. The predicted model was further optimized by molecular dynamics simulation

**Table 1.**
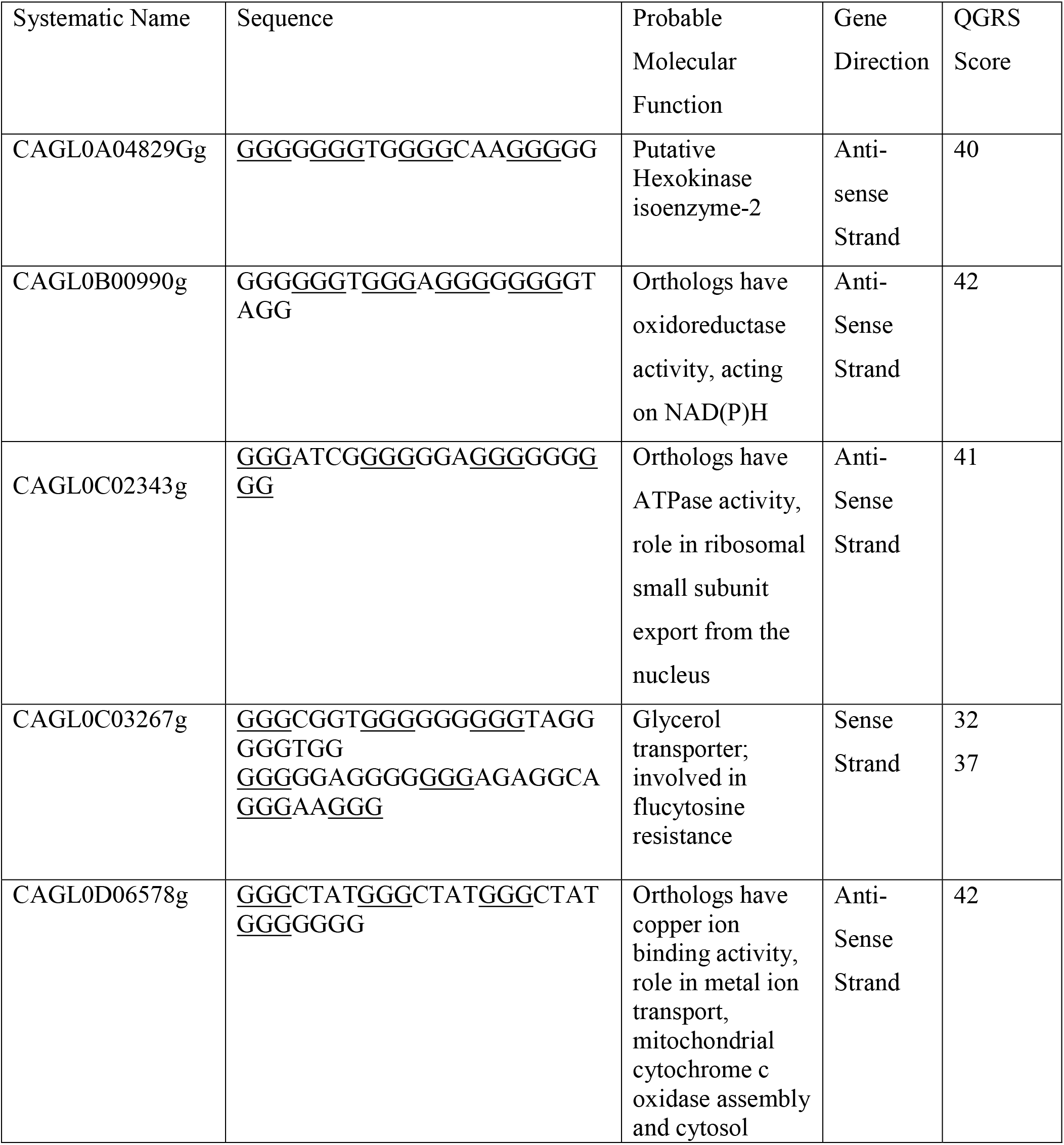
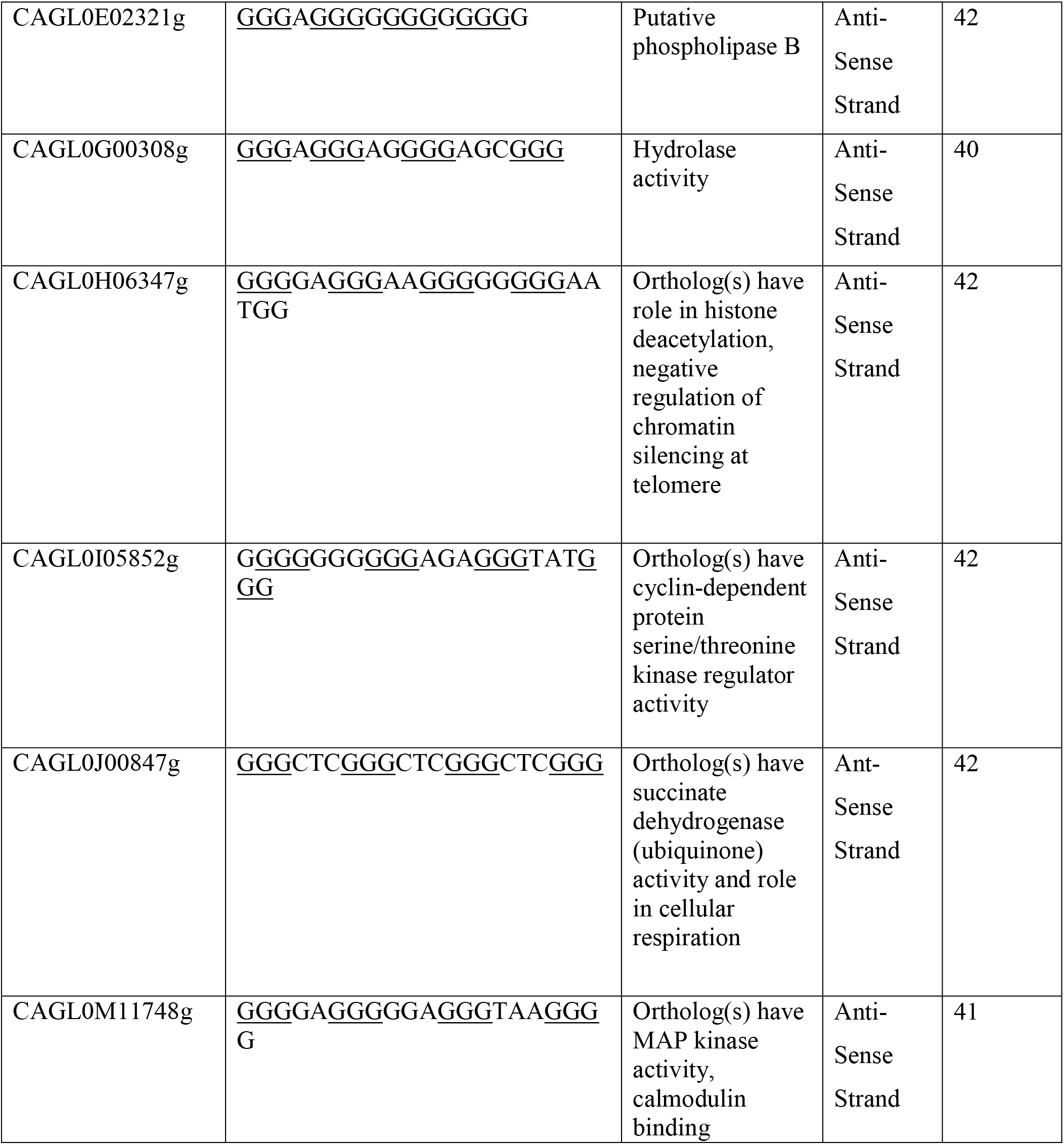
depicted the details of the G-quadruplex forming sequence occurring with more than 98% frequency and present in the crucial genes of chromosome A-M of the strains of *Bg2 Candida*.

### 2.3 Structure Designing and NMR Studies of bis(1,3,7-trimethyl-2,6-dioxo-2,3,6,7,8,9-hexahydro-1H-purin-8-yl)gold (Compound A), bis(1,3,7,9-tetramethyl-2,6-dioxo-2,3,6,7,8,9-hexahydro-1H-purin-8-yl)gold (Compound B) and bis(3,7,9-trimethyl-2,6-dioxo-2,3,6,7,8,9-hexahydro-1H-purin-8-yl)gold (Compound C)

The structure of Compound A, B, and C was designed using MarvinSketch Software based on the following parameter: Modification can occur at the N9 position of the Xanthine derivative conjugated with Gold in the scaffold causes loss of G-quadruplex binding selectivity and potency. Modification occurs at the N1 position of Xanthine derivative conjugated with Gold in the scaffold can retain the planer geometry of the designed structure (Meier-Menches et al., 2020). ^1^H and ^13^C NMR spectra of compounds A, B, and C were determined using an NMR prediction plugin with MarvinSketch Tool. The NMR spectra of compounds A, B, and C were recorded in DMSO-d_6_ [deuterated dimethyl sulfoxide]. The tetramethylsilane was used as the reference sample, and the frequency was adjusted at 500 MHz.

### 2.4 Molecular Docking Studies

The proposed 3D model of the SDH1 G-quadruplex structure was used as a molecular target for docking studies. Both the proposed SDH1 G-quadruplex structure and ligands were prepared using the Autodock tool [Morris et al., 2009]. After computing the Gasteiger charges and polar hydrogen bonds, docking studies were done with the AutoDock4.2 program using the Lamarckian genetic algorithm [Morris et al., 2009]. The Grid box size was constrained to (100 ×100 ×100) A° along the axes (x,y, z), with a grid spacing of 0.375 A°. For ligands, 25 GA runs were performed with a population size of 150 random individuals. For each ligand, the maximum number of the evaluation is set to 2.5×10^7^ The rate of gene mutation and cross-over is set to 0.02 and 0.8, respectively. In docking simulation, ligands were allowed full flexibility, whereas the SDH1 G-quadruplex structure was not fully flexible and kept rigid. The best conformers were chosen based on the binding free energy.

### 2.5 ADMET analysis of compounds A, B, and C

Pharmacokinetics properties like absorption, distribution, metabolism, excretion, and toxicity parameters of compounds A, B, and C were analyzed using the pkCSM webserver. The pkCSM webserver determined ADMET profiling based on the ADMET descriptor algorithm (Pires et al., 2015). ADMET profiling allows understanding the drug candidate’s safety and efficiency and deciphers the viability of the drug [Guan et al., 2019]. This ADMET study includes drug-like properties, lipophilicity, pKa, the volume of distribution, absorption, transporter, solubility, metabolism, hepatic clearance, etc. The absorption of the compounds correlates with the factor, including permeability through the membrane (specify by Caco-2 cell line), P-glycoprotein substrate, or inhibitor. The absorption also depends on the skin permeability and absorption through the intestine. The distribution of the compound (Drug to be molecule) correlates with the factors such as blood-Brain-Barrier, permeability through CNS. The distribution also depends on the volume of the distribution of the drug molecule. Metabolism of the drug molecule depends on the CYP substrate and inhibitor models such as CYP3A4, CYP2C19, CYP2D6, CYP2D6, CYP1A2, and CYP3A4. The excretion of the compound is predicted by renal OCT2 substrate and the total clearance model. In comparison, the drug’s toxicity is determined based on the hERG inhibition, AMES toxicity, and hepatotoxicity.

## 3. Result and Discussion

### 3.1 Putative G-quadruplex forming sequences in *Candida glabrata* genome

Presently, there are less effective antifungal agents available to treat fungal infections caused by *Candida* species. The emergence of latent disease and antifungal drug-resistant strains rings a worldwide alarm to identify novel antifungal drug targets. Depending on the nucleotide sequence, the nucleic acid may acquire various distinct secondary structures, including duplex, triplex, and G-quadruplex (Burge et al., 2006). Among all putative secondary structures, G-Quadruplex has been depicted as the utmost studied evolutionary conserved drug target. An array of smallmolecule compounds have been developed to regulate G-Quadruplex structures by stabilizing or destabilizing and may be used as a promising drug target in the battle against fungal infections.

The genomic sequence of *Candida glabrata* was extracted from the NCBI database and searched for the presence of PQS by utilizing a “Putative G-quadruplex Prediction Tool algorithm” developed by the Indian Institute of Technology, Indore (Mishra et al., 2016). All predicted G-quadruplex sequences were then confirmed by QGRS Mapper (Kikin et al., 2006) and PQSFinder (Hon et al., 2017). AS G-quadruplex folding solely depends on consecutive G-tetrad loops and different patterns. The constraint has been set to 3 or 4 nucleotides as the minimum length in G-tetrad and 0 and 5 nucleotides as the minimum and maximum length of the loop. The G-quadruplex prediction tool with the above set parameters was utilized on the *Candida glabrata* strains, identifying more than 5000 putative G-quadruplex forming sequences. But selected only those sequences that are present in genes play a crucial role in virulence and pathogenesis. PQS present in essential genes is shown in table 1 and supplementary table 1.

Considering the likeness between two sequences may correlate with similar functions. For conquering multidrug-resistant fungal infection, evolutionary conservation is essential that satisfy one of the vital criteria to work as a promising drug target. Therefore we utilized the BLAST and GO annotation to determine the probability of each cluster in *Candida glabrata*.

### 3.2 Proposed 2-D and 3-D model of SDH1 G-Quadruplex

The secondary structure of the G-quadruplex forming sequence in the SDH1 gene of *Candida glabrata* was predicted using the RNAfold web server shown in figure 1, and its stability was found to be ΔG°: −19.6 Kcal/mol. The entropy vs. position graph (figure 1) shows that the trinucleotide (CUC) loop has more entropy, whereas G-tetrad’s forming guanine nucleotide have less entropy. Figure 10 Depicts the hairpin conformation of SDH1 PQS, and the graph represents each nucleotide’s positional entropy. Since the structure of SDH1 G-quadruplex is not modeled, so we obtained a predicted model on the basis of parallel G4 that is telomeric G4 AGGG[TTAGGG]_3_ {PDB ID-4G0F}. The SDH1 G-quadruplex structure consists of three G-tetrads (G2-G8-G14-G20; G3-G9-G15-G21; G4-G10-G16-G22) interconnected by three CUC loops. G2-G8-G14-G20 guanine nucleotides were interconnected by hoogsteen base-pairing where N7, O6 of G2 formed hydrogen bonds with N2 and N1 of G20 nucleotide, N7, O6 of G20 formed hydrogen bond with N1 and N2 of G14, whereas N7, O6 of G20 formed hydrogen bond with N1 and N2 of G8. All these G2-G8-G14-G20 together formed G-tetrad and stabilized by K^+^ ion where K^+^ formed a bond with O6 of all G’s of the tetrad, which help stabilize the G-tetrad.

**Figure 1:**
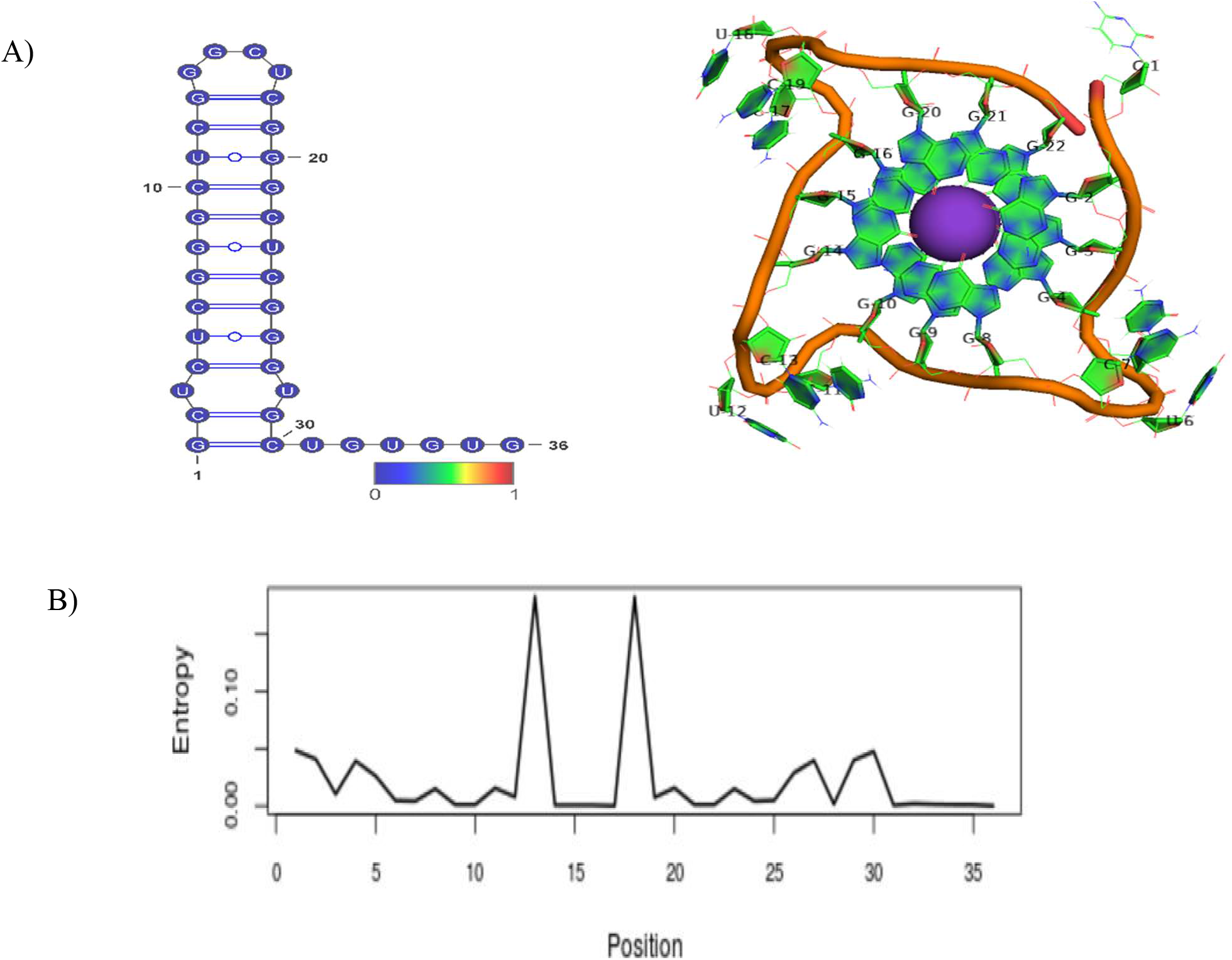
**A)** Proposed hairpin conformation having ΔG°: −19.6 Kcal/mol and three-dimensional G4 structure of PQS present in SDH1 gene, B) Positional entropy for each nucleotide

### 3.3 In-silico Characterization of Compounds A, B, and C Using MarvinSketch NMR tool

#### 3.3.1 Compound A (bis(1,3,7-trimethyl-2,6-dioxo-2,3,6,7,8,9-hexahydro-1H-purin-8-yl)gold)

DMSO-d_6_ [deuterated dimethyl sulfoxide] was used as a solvent for the in-silico characterization of compound A. At the same time, Trimethylsilane was utilized as a reference sample, and NMR spectra were recorded at 500 MHz frequency

^1^H NMR (500 MHz, DMSO-d_6_) *δ* 2.93 (s, 3H, CH_3_ (Atom no- 14,14,14,28,28,28)), 3.43 (s, 3H, CH_3_ (Atom no- 8,8,8,22,22,22)), 3.91 (s, 3H, CH_3_ (Atom no- 10,10,10,24,24,24)), 7.15 (s, 1H, CH (Atom no- 31,33)), 9.15 (s, 1H, NH (Atom no- 30,32). ^1^H NMR spectrum shown in figure 2a

**Figure 2a:**
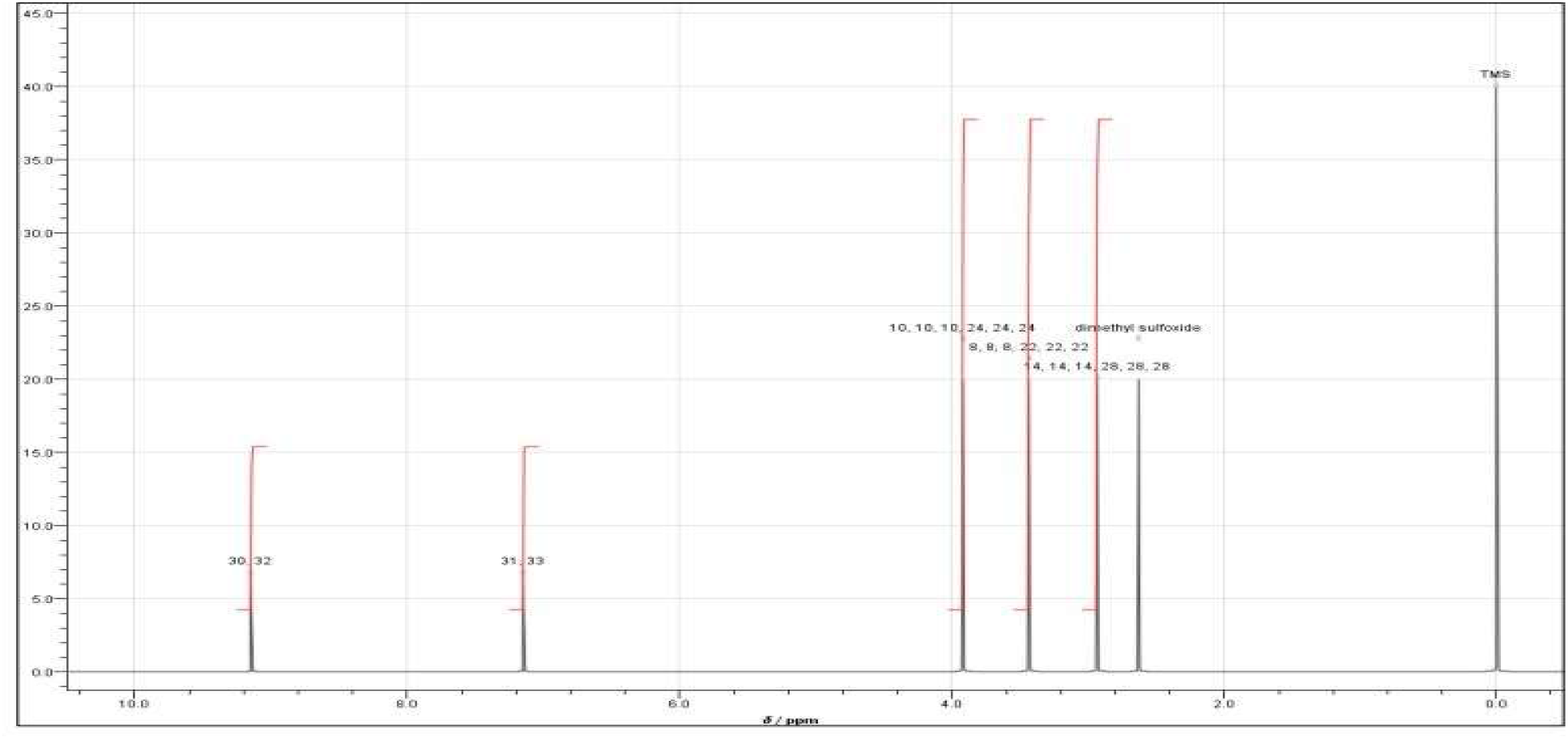
This figure depicts ^1^H NMR spectrum of (bis(1,3,7-trimethyl-2,6-dioxo-2,3,6,7,8,9-hexahydro-1 H-purin-8-yl)gold)

^13^C NMR (500 MHz, DMSO-d6) *δ* 28.20 (s, CH_3_ (Atom no- 8,22)), 29.90 (s, CH_3_ (Atom no- 14,28)), 32.14 (s, CH_3_ (Atom no- 10,24)), 65.69 (s, CH (Atom no- 12,26)), 134.21 (s, C-N (Atom no- 5,19)), 151.90 (s, CO (Atom no- 2,16)), 153.09 (s, C-N (Atom no- 4,18)), 156.43 (s, CO (Atom no- 6,20))..

#### 3.3.2 Compound B (bis(1,3,7,9-tetramethyl-2,6-dioxo-2,3,6,7,8,9-hexahydro-1H-purin-8-yl)gold)

DMSO-d_6_ [deuterated dimethyl sulfoxide] was used as a solvent for the in-silico characterization of compound B. At the same time, Trimethylsilane was utilized as a reference sample, and NMR spectra were recorded at 500 MHz frequency.

^1^H NMR (500 MHz, DMSO-d6) *δ* 2.92 (s, 3H, CH_3_ (Atom no- 30,30,30,31,31,31)), 2.94 (s, 3H, CH_3_ (Atom no- 14,14,14,28,28,28)), 3.43 (s, 3H, CH_3_ (Atom no- 8,8,8,22,22,22)), 3.96 (s, 3H, CH_3_ (Atom no- 10,10,10,24,24,24)), 7.15 (s, 1H, CH (Atom no- 32,33). ^1^H NMR spectrum shown in figure 2b.

**Figure 2b:**
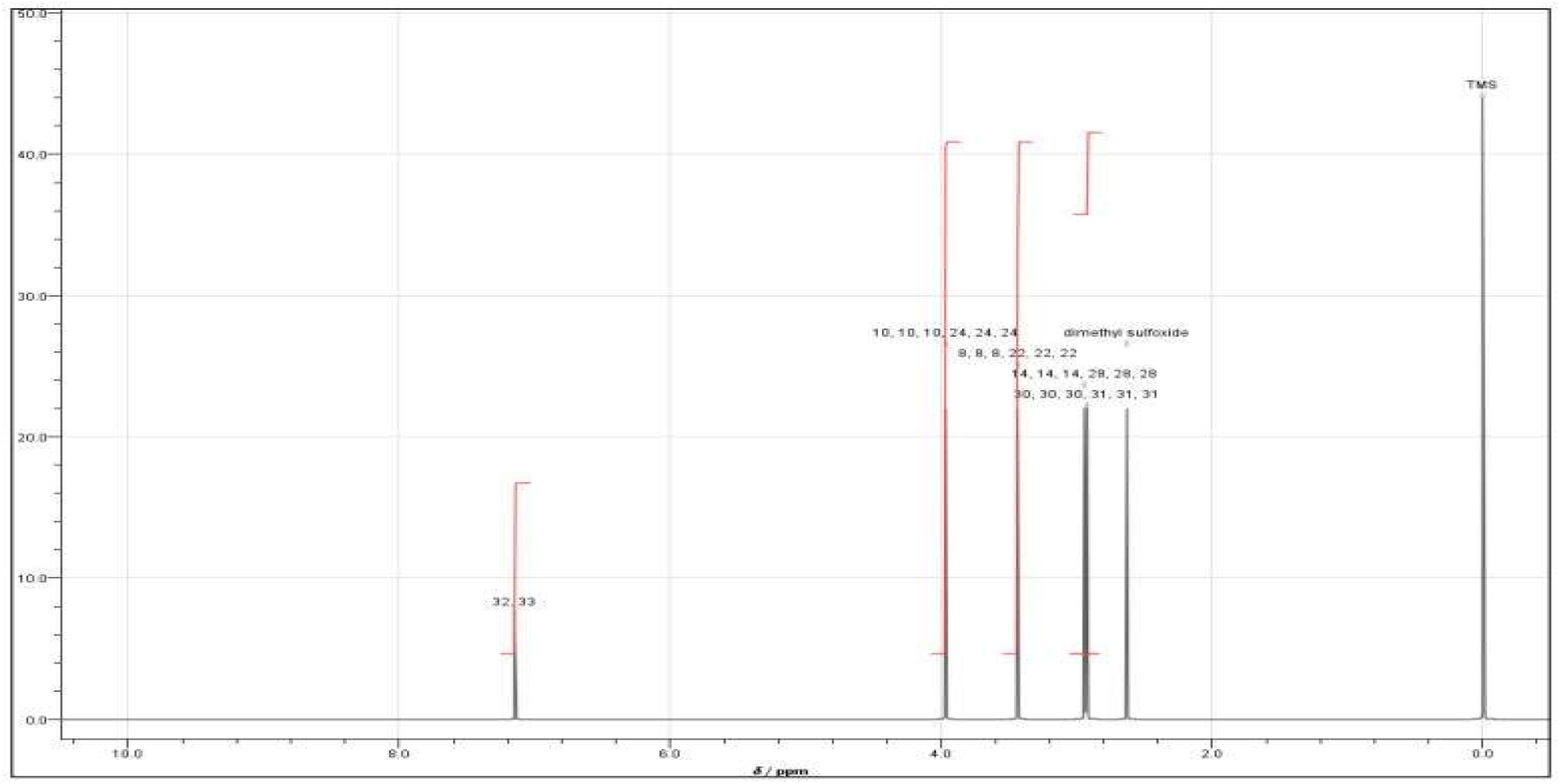
This figure depicts ^1^H NMR spectrum of (bis(1,3,7,9-tetramethyl-2,6-dioxo-2,3,6,7,8,9-hexahydro-1H-purin-8-yl)gold)

^13^C NMR (500 MHz, DMSO-d6) *δ* 28.20 (s, CH_3_ (Atom no- 8,22)), 30.19 (s, CH_3_ (Atom no- 14,28)), 32.14 (s, CH_3_ (Atom no- 10,24)), 34.72 (s, CH_3_ (Atom no- 30,31)), 68.66 (s, CH (Atom no- 12,26)), 138.54 (s, C-N (Atom no- 5,19)), 151.90 (s, CO (Atom no- 2,16)), 156.42 (s, C-N (Atom no- 4,18)), 156.27 (s, CO (Atom no- 6,20)).

#### 3.3.3 Compound C (bis(3,7,9-trimethyl-2,6-dioxo-2,3,6,7,8,9-hexahydro-1H-purin-8-yl)gold)

DMSO-d_6_ [deuterated dimethyl sulfoxide] was used as a solvent for the in-silico characterization of compound C. At the same time, Trimethylsilane was utilized as a reference sample, and NMR spectra were recorded at 500 MHz frequency.

^1^H NMR (500 MHz, DMSO-d6) *δ* 2.95 (s, 3H, CH_3_ (Atom no- 28,28,28,29,29,29)), 2.96 (s, 3H, CH_3_ (Atom no- 13,13,13,26,26,26)), 4.03 (s, 3H, CH_3_ (Atom no- 9,9,9,22,22,22)), 7.20 (s, 1H, CH (Atom no- 30,31), 9.76 (s, 1H, NH (Atom no- 1,19). ^1^H NMR spectrum shown in figure 2c.

**Figure 2c:**
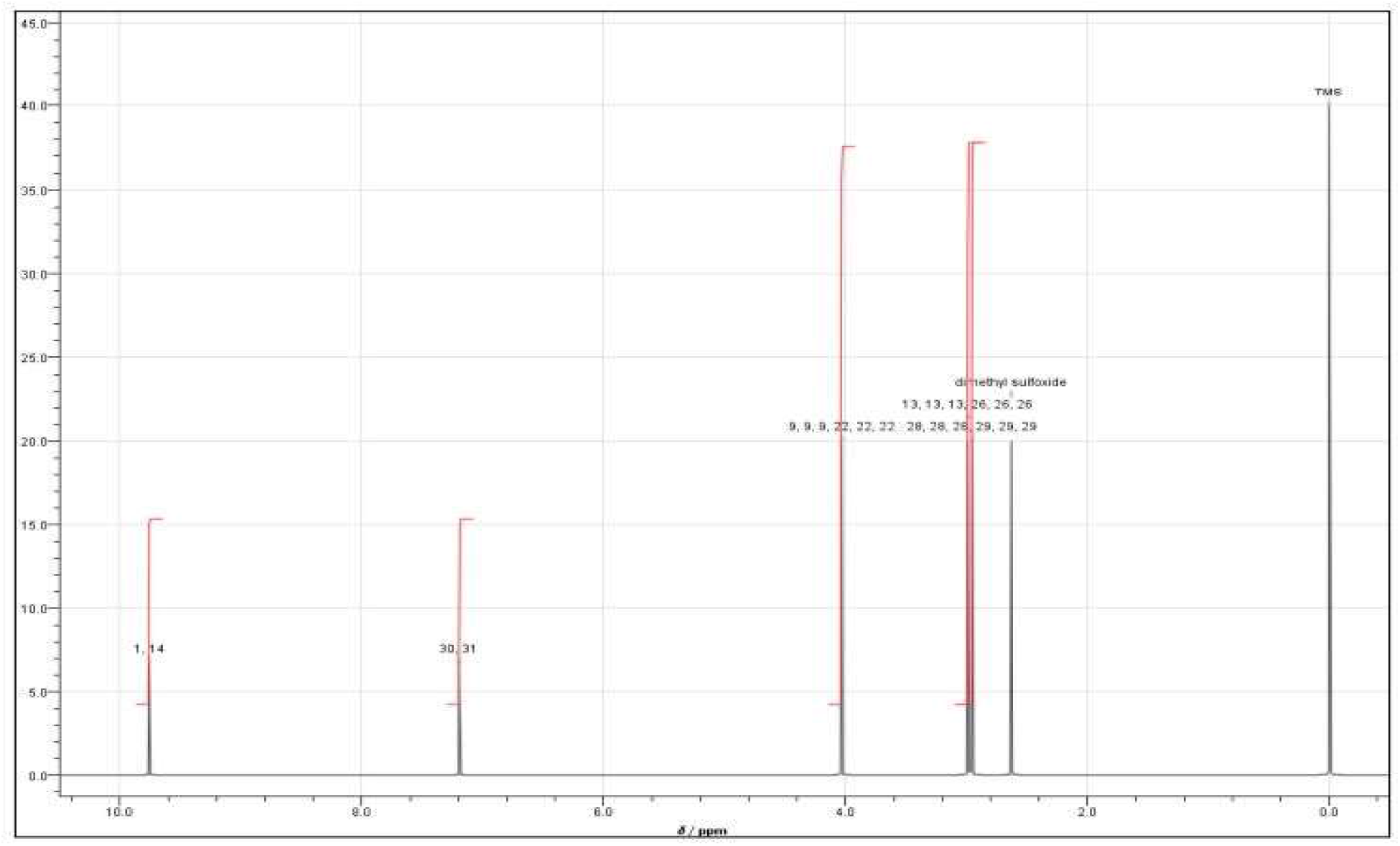
This figure depicts ^1^H NMR spectrum of (bis(3,7,9-trimethyl-2,6-dioxo-2,3,6,7,8,9-hexahydro-1 H-purin-8-yl)gold)

^13^C NMR (500 MHz, DMSO-d6) *δ* 29.76 (s, CH_3_ (Atom no- 13,26)), 32.14 (s, CH_3_ (Atom no- 9,22)), 34.84 (s, CH_3_ (Atom no- 28,29)), 68.97 (s, CH (Atom no- 11,24)), 135.51 (s, C-N (Atom no- 5,18)), 151.50 (s, CO (Atom no- 2,15)), 156.64 (s, CO (Atom no- 6,19)), 158.62 (s, C-N (Atom no- 4,17)).

### 3.4 In-silico ADMET Studies for Compound A, B, and C

In-silico ADMET study was used to depict various pharmacokinetic parameters shown in table 2. It is suggested as an initial step to analyse chemical compound entity (novel drug) to prevent resources and time on the lead compound that the human body may metabolize in an inactive form or maybe cause toxicity and unable to penetrate the membrane. The calculated LogP (Octonal-water diffusion coefficient) for Compound A, B, and C was 0.0, which shows compounds A, B, and C was slightly lipophilic, whereas LogS indicated solubility. LogS value for compounds A, B, and C was found to be −4.619 log Mol/L, −4.905 log Mol/L, and −3.961 log Mol/L. These values of LogS indicated that compounds A, B, and C are very soluble. In the case of Intestinal absorption, the calculated intestinal absorption for compound A was 60.918%, for compound B, it was 67.487%, and compound C has 53.185% absorption through the intestine. The result shows that compound B has the highest intestinal absorption, followed by compound A and compound C at least intestinal absorption. Higher intestinal absorption shows compounds A, B, and C could be adequately absorbed upon oral administration. The compounds A, B, and C were analyzed for skin permeability, and the calculated skin permeability for compound A was −2.917 log K_p_, compound B was −2.911 log K_p_, and compound C was −2.769 log K_p_. All the compounds have skin permeability log K_p_ < −2.5, which shows that compounds A has higher skin permeability than the other two compounds. In predicting the substrate/inhibitor for P-glycoprotein, Compound A was found to be non-inhibitor and non-substrate. Compound B comes out to be the substrate for P-glycoprotein and non-inhibitor of P-glycoprotein.

**Table 2:**
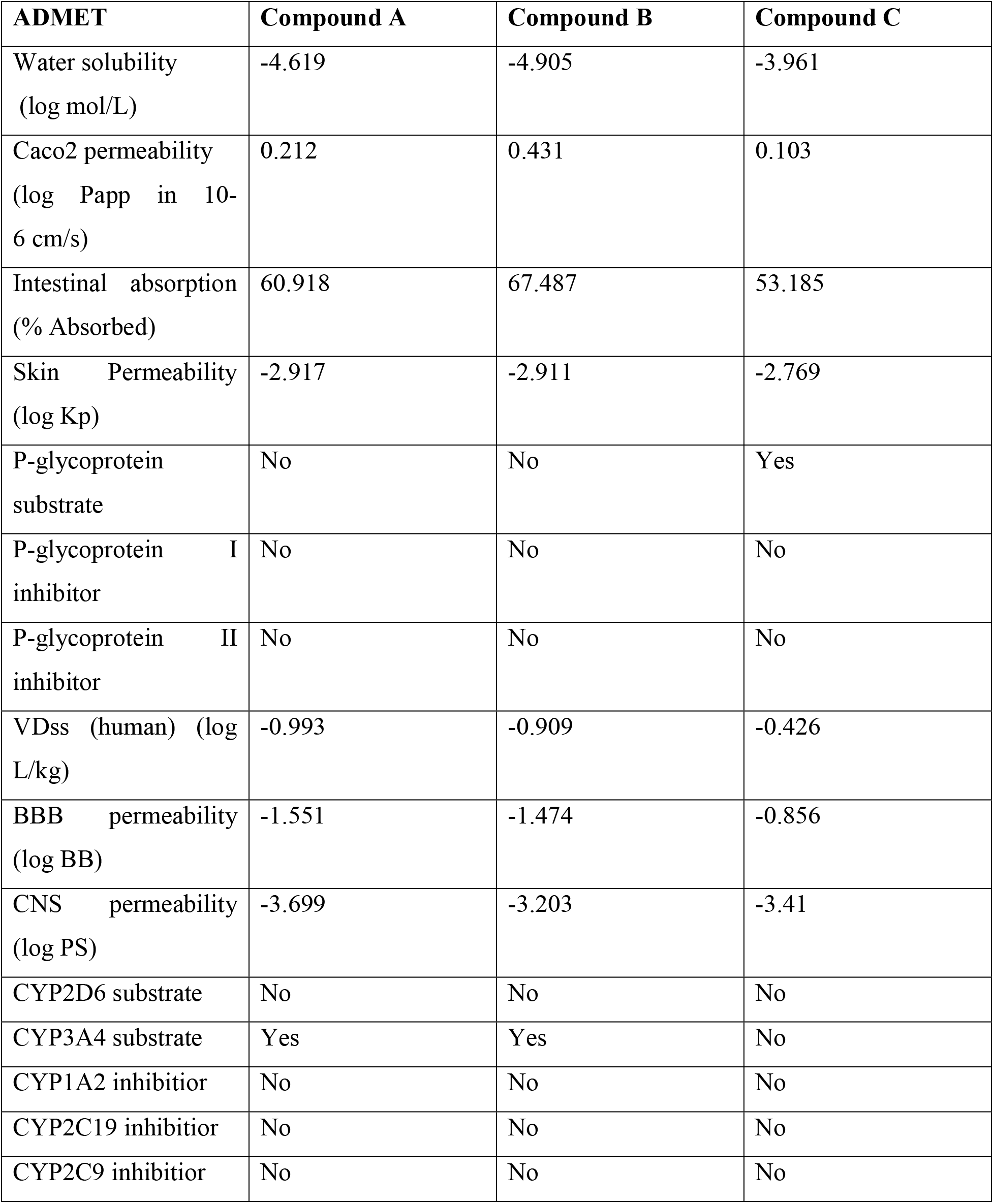
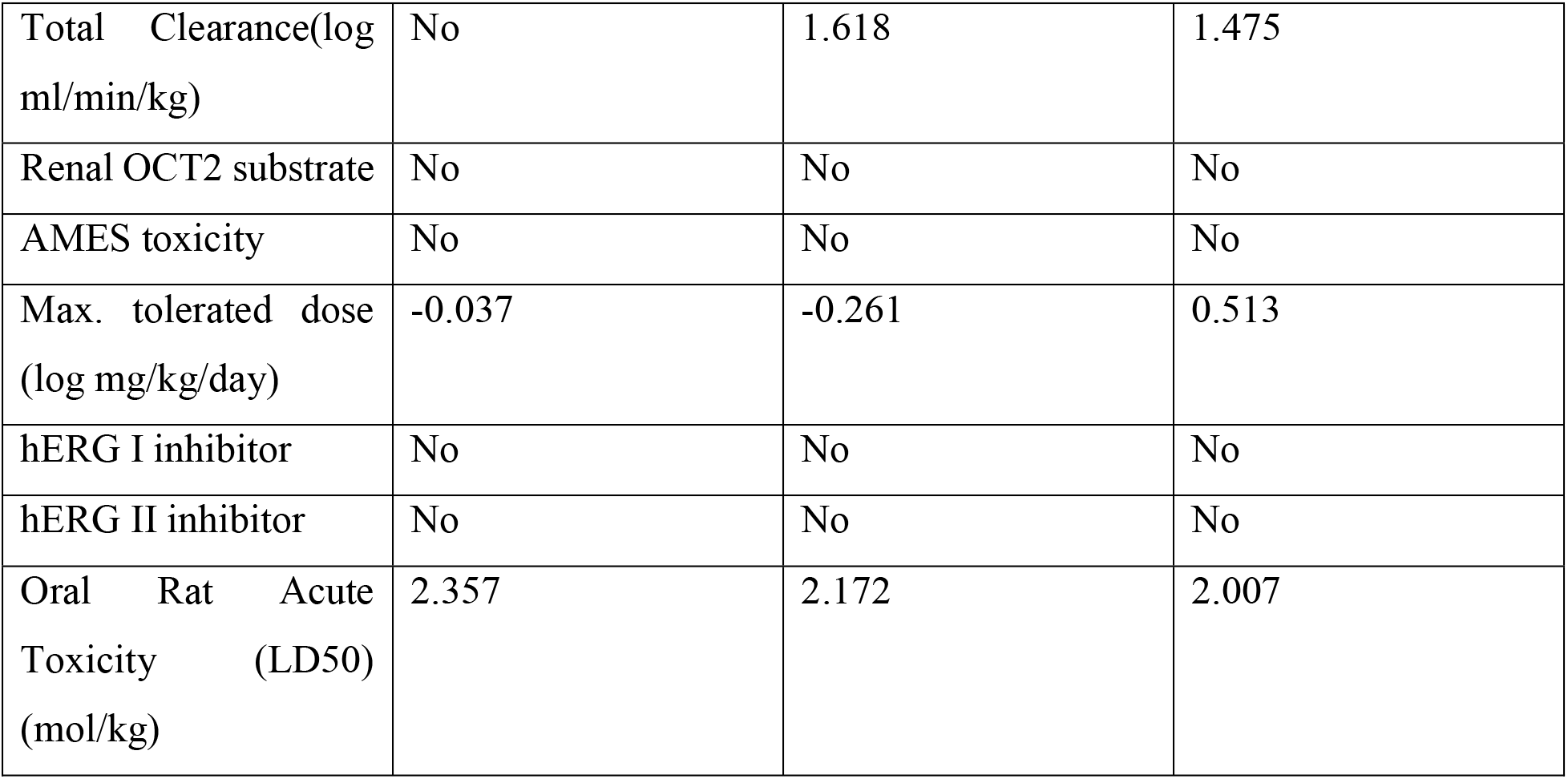
ADMET analysis of Compound A, B, and C:

| **ADMET** | **Compound A** | **Compound B** | **Compound C** |
| --- | --- | --- | --- |
| Water solubility<br>(log mol/L) | -4.619 | -4.905 | -3.961 |
| Caco2 permeability<br>(log Papp in 10-6 cm/s) | 0.212 | 0.431 | 0.103 |
| Intestinal absorption<br>(% Absorbed) | 60.918 | 67.487 | 53.185 |
| Skin Permeability<br>(log Kp) | -2.917 | -2.911 | -2.769 |
| P-glycoprotein<br>substrate | No | No | Yes |
| P-glycoprotein I<br>inhibitor | No | No | No |
| P-glycoprotein II<br>inhibitor | No | No | No |
| VDss (human) (log<br>L/kg) | -0.993 | -0.909 | -0.426 |
| BBB permeability<br>(log BB) | -1.551 | -1.474 | -0.856 |
| CNS permeability<br>(log PS) | -3.699 | -3.203 | -3.41 |
| CYP2D6 substrate | No | No | No |
| CYP3A4 substrate | Yes | Yes | No |
| CYP1A2 inhibitor | No | No | No |
| CYP2C19 inhibitor | No | No | No |
| CYP2C9 inhibitor | No | No | No |
| Total Clearance(log ml/min/kg) | No | 1.618 | 1.475 |
| Renal OCT2 substrate | No | No | No |
| AMES toxicity | No | No | No |
| Max. tolerated dose (log mg/kg/day) | -0.037 | -0.261 | 0.513 |
| hERG I inhibitor | No | No | No |
| hERG II inhibitor | No | No | No |
| Oral Rat Acute Toxicity (LD50) (mol/kg) | 2.357 | 2.172 | 2.007 |

In comparison, compound C was found to be a non-substrate and non-inhibitor of P-glycoprotein. The calculated value for Blood-Brain-Barrier permeability for compounds A, B, and C comes out low and unable to cross the BBB. The main CYPs used in metabolism were CYP1A2, CYP2C9, CYP2C19, and CYP3A4, and compounds A, B, and C were found to be non-inhibitory and nonsubstrate for CYP1A2, CYP2C9, CYP2C19, and CYP3A4. AMES Toxicity was exploited to analyze whether these compounds were mutagenic or not. It is found that compounds A, B, and C were non-mutagenic as well as non-carcinogenic. Total clearance in terms of excretion for compound A was 1.508 log ml/min/kg. Compound B has a total clearance of 1.618 log ml/min/kg, and compound C has 1.475 log ml/min/kg. The calculated oral rat toxicity (LD50) for compound A was found to be 2.357 mol/kg, LD50 for compound B comes out 2.172 mol/kg, and for compound C, it came out to be 2.007 mol/kg.

### 3.5 Molecular Docking Analysis

Molecular docking has been performed using AutoDock software, and the best binding pose based on the lowest free energy was chosen. The molecular interaction between SDH1 G4-compound complexes was analyzed using discovery studio software shown in table 3. Compound A was docked between the G-tetrads and C_5_U_6_C_7_ loop shown in figure 3. The C_7_ is present in the loop, and G_15_ of G-tetrad forms a carbon-hydrogen bond with compound A, respectively shown in light green color, and G_14_ and G_20_ of G-tetrads binds with compound A through hydrogen bond shown in green color and. Whereas G_20_ also formed metal-acceptor bond with a gold atom present in the center of the compound A and Pi-Pi stacking interaction and a Pi-alkyl bond with the compound A. The estimated inhibition constant for compound A was calculated to be 1.73 μM. In Compound B (figure 4), C_7_ present in the loop of SDH1 G4 binds to compound B through the carbon-hydrogen bond and conventional hydrogen bond. At the same time, G_4_ and G_10_ of G-tetrads form a carbon-hydrogen bond with compound B. G_4_, and G_10_ also participated in the Pi-Pi stacking, and G_4_, G_8_, G_10_ includes Pi-alkyl interaction with compound B. The estimated inhibition constant was found to be 15.62 μM. In terms of compound C (figure 5), C_13_ and C_11_ present in the loop of SDH1 G4 structure bind with the compound C through carbon-hydrogen bond and conventional hydrogen bond, respectively. G_10_ and G_14_ of G-tetrad participate in the carbon-hydrogen bond, and G_4_ also formed a metal-acceptor bond with compound C. At the same time, G_15_ and G_10_ formed Pi-Anion and Pi-Sigma with compound C, respectively, whereas G_10_ formed Pi-Pi-stacked and Pi-Alkyl bonds with compound C. The estimated inhibition constant for compound C was calculated to be 24.37 μM.

**Table 3:**
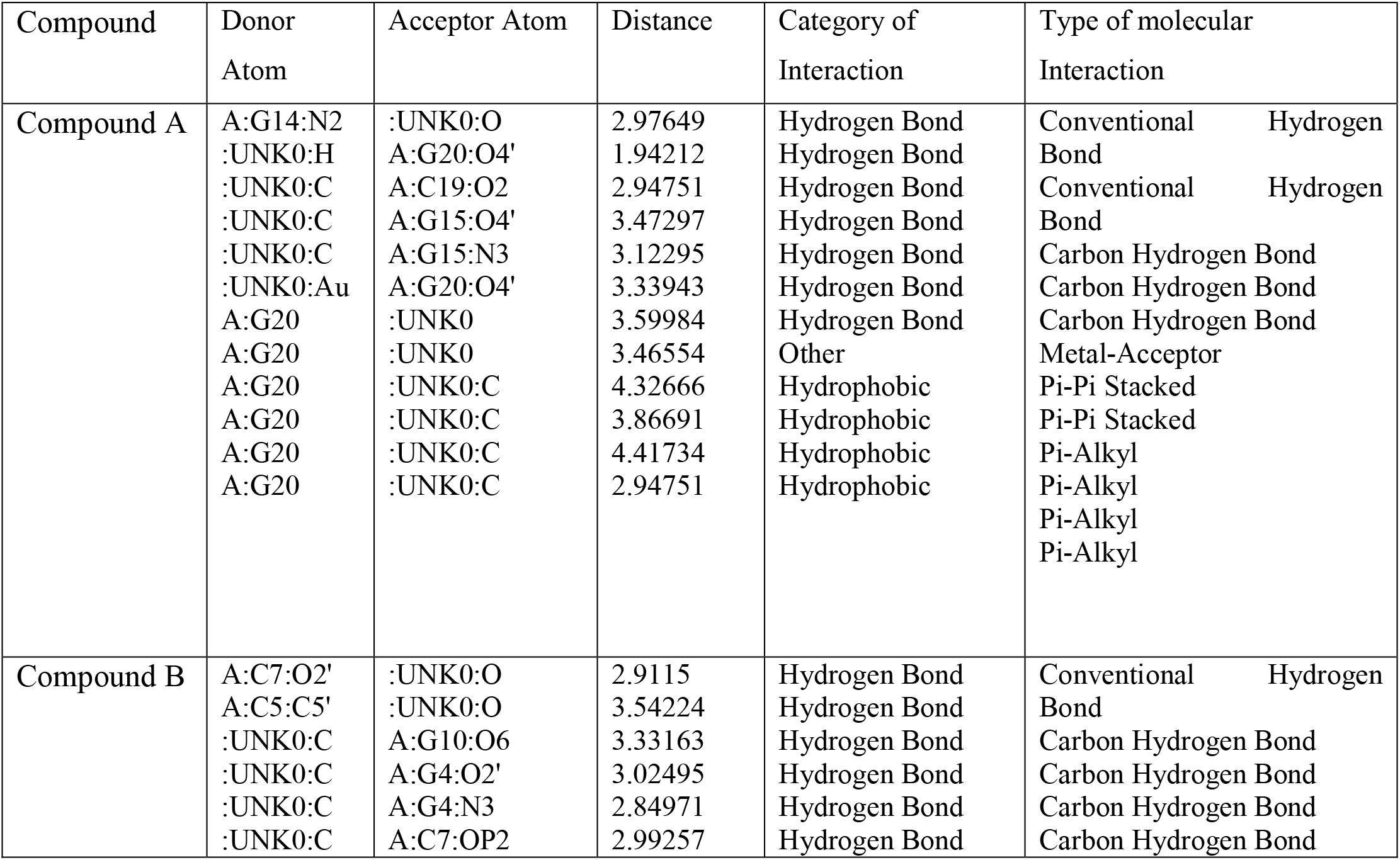
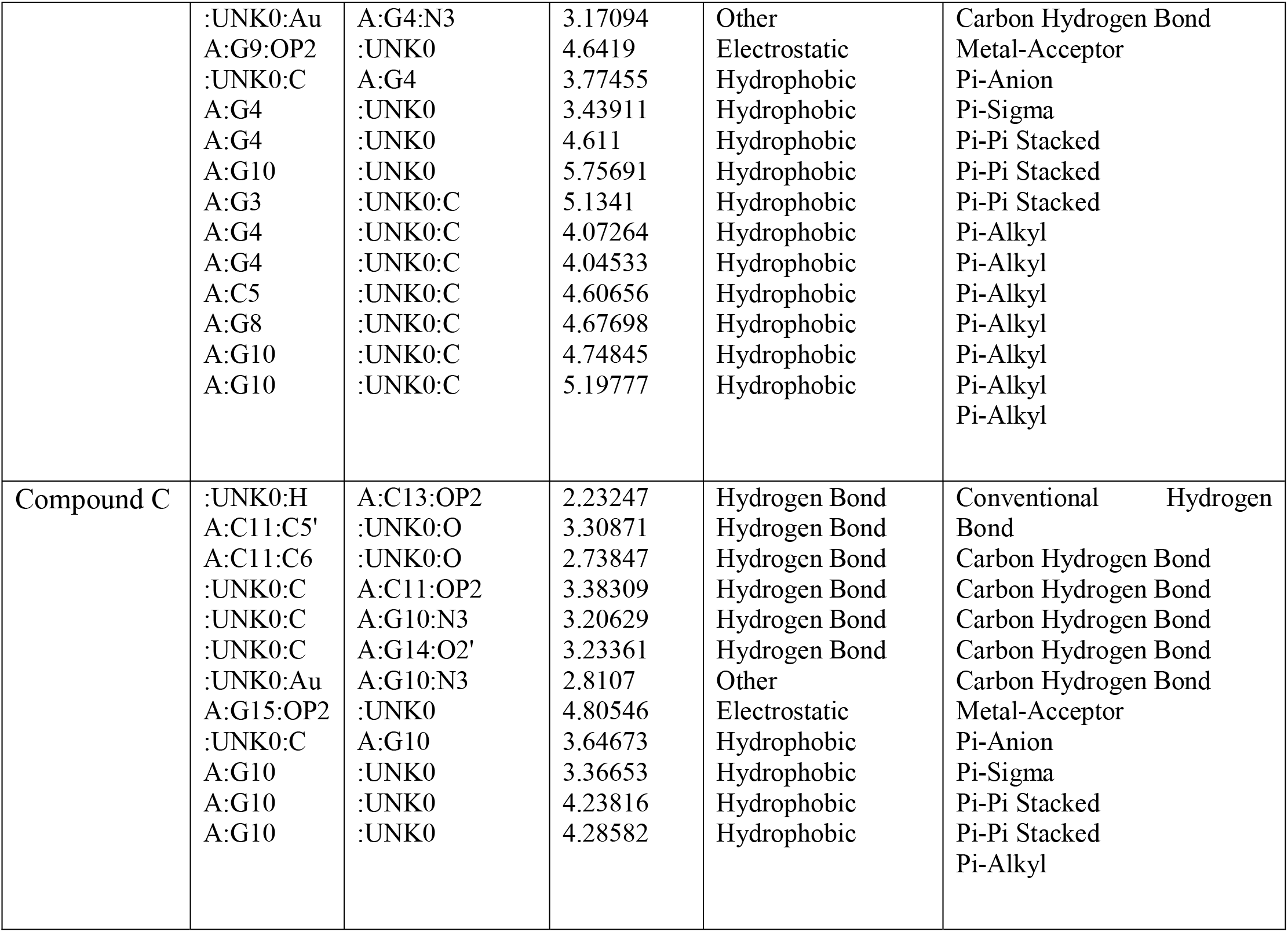
The molecular interaction of G4 structure of PQS in SDH1 and Compounds A, B, and C

**Figure 3:**
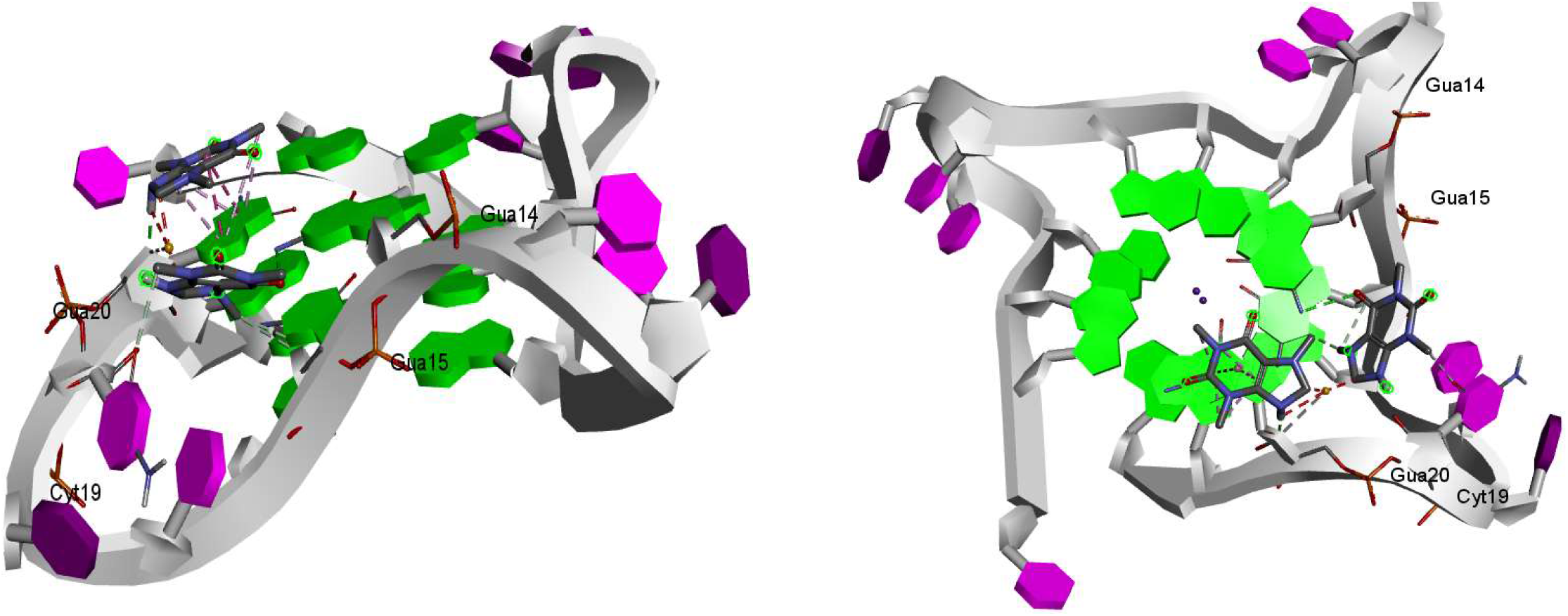
Molecular interaction representation of compound A with G4 structure of PQS in SDH1 via side view and top view

**Figure 4:**
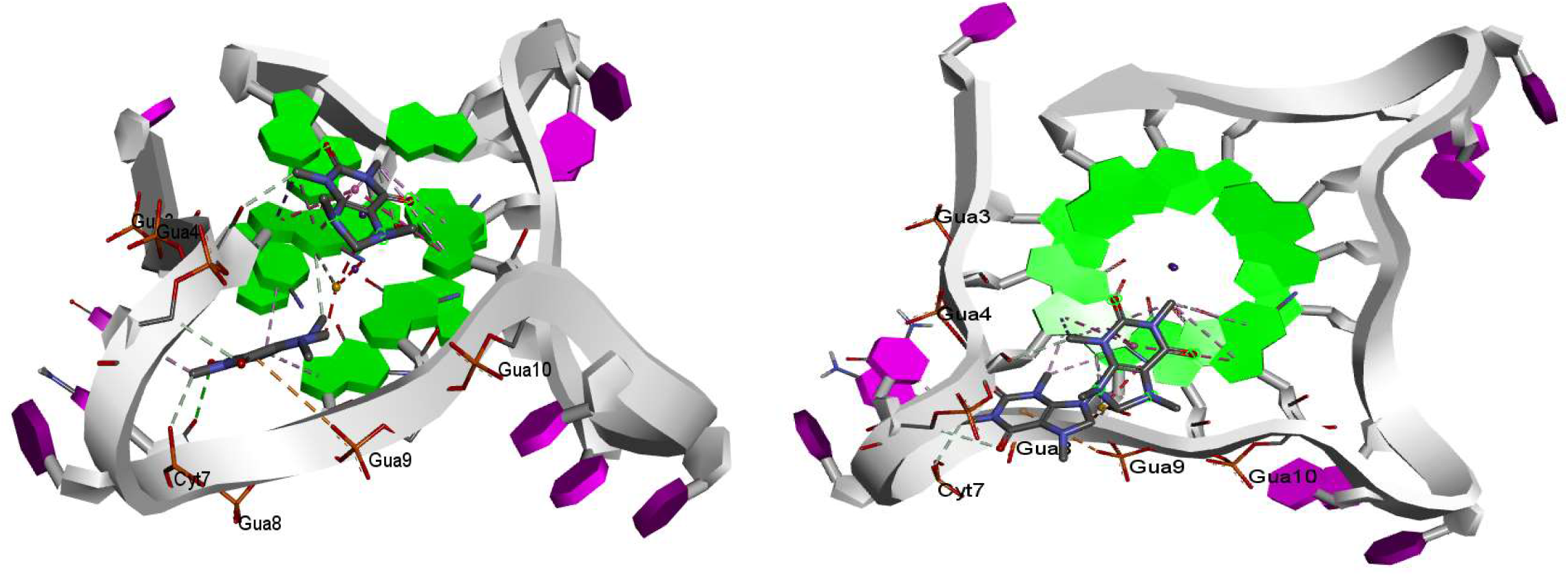
Molecular interaction representation of compound B with G4 structure of PQS in SDH1via side view and top view

**Figure 5:**
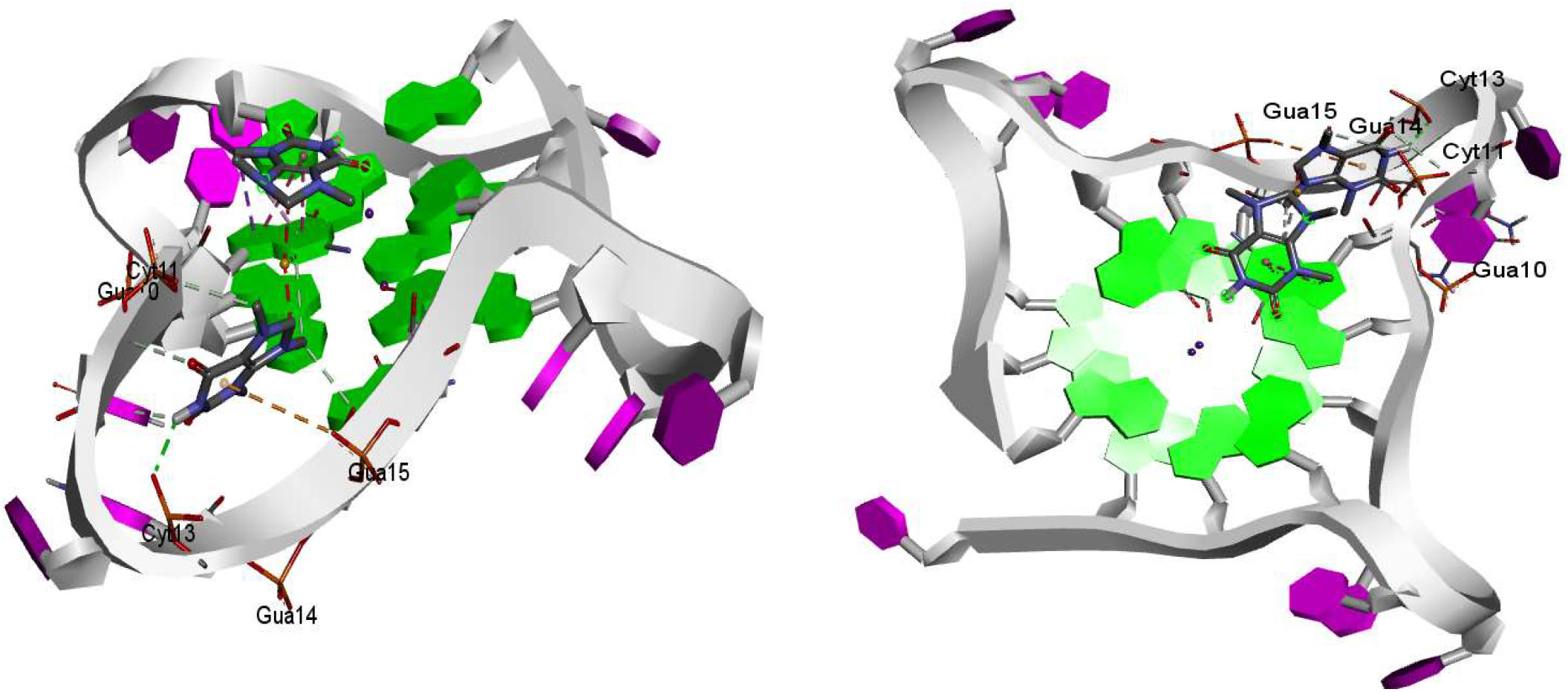
Molecular interaction representation of compound C with G4 structure of PQS in SDH1via side view and top view

## 4. Conclusion

Determining the putative quadruplex sequences within the pathogenic or drug resistance genes of *Candida glabrata* has a focussed structure that can be targeted by G-quadruplex binding drugs such as Gold Carbene and their derivatives to inhibit *Candida glabrata* pathogenicity. We have to determine whether these G4 structures are formed and may modulate biological and cellular functions. Thus, targeting PQS in *Candida glabrata* could show a novel antifungal target to ameliorate *Candida glabrata* infection. For this, we have searched for the presence of PQS by utilizing a “Putative G-quadruplex Prediction Tool algorithm”. QGRS Mapper and PQSFinder then confirmed all predicted G-quadruplex sequences. Afterward, We have proposed a G4 structure for PQS present in the SDH1 from all predicted PQS. It may be a key target to ameliorate the *C.glabrata* infection because it encodes a protein that plays a vital role in energy production in *C.glabrata* cells shown in the figure. The structure was built based on the already current structure that is telomeric G4 AGGG (TTAGGG)_3_ {PDB ID-4G0F} for identifying its stabilization by Gold carbine derivatives. Molecular docking and ADMET analysis show that Compound A has the highest binding affinity and has the best ADMET properties among the two compounds. The present study represents PQS in the SDH1 gene could be a novel antifungal target. Thus these three compounds may be an effective therapeutic option for infections related to Candida glabrata. In the future, these experimental and computational should be correlated with biophysical techniques to ascertain the validity of these data.

## Reference

Pappas, P. G., Lionakis, M. S., Arendrup, M. C., Ostrosky-Zeichner, L., & Kullberg, B. J. (2018). Invasive candidiasis. Nature Reviews Disease Primers, 4(1), 18026. https://doi.org/10.1038/nrdp.2018.26

Fisher, M. C., Hawkins, N. J., Sanglard, D., & Gurr, S. J. (2018). Worldwide emergence of resistance to antifungal drugs challenges human health and food security. Science (New York, N.Y.), 360(6390), 739–742. https://doi.org/10.1126/science.aap7999

Vautier, S., MacCallum, D. M., & Brown, G. D. (2012). C-type lectin receptors and cytokines in fungal immunity. Cytokine, 58(1), 89–99. https://doi.org/10.1016/j.cyto.2011.08.031

Gow, N. A. R., Latge, J.-P., & Munro, C. A. (2017). The Fungal Cell Wall: Structure, Biosynthesis, and Function. Microbiology Spectrum, 5(3). https://doi.org/10.1128/microbiolspec.FUNK-0035-2016

Kumamoto, C. A. (2011). Inflammation and gastrointestinal Candida colonization. Current Opinion in Microbiology, 14(4), 386–391. https://doi.org/10.1016/j.mib.2011.07.015

Vincent Vincent, J.-L., Rello, J., Marshall, J., Silva, E., Anzueto, A., Martin, C. D., Moreno, R., Lipman, J., Gomersall, C., Sakr, Y., Reinhart, K., & EPIC II Group of Investigators. (2009). International study of the prevalence and outcomes of infection in intensive care units. JAMA, 302(21), 2323–2329. https://doi.org/10.1001/jama.2009.1754

Turner, S. A., & Butler, G. (2014). The Candida pathogenic species complex. Cold Spring Harbor Perspectives in Medicine, 4(9), a019778. https://doi.org/10.1101/cshperspect.a019778

Dadar, M., Tiwari, R., Karthik, K., Chakraborty, S., Shahali, Y., & Dhama, K. (2018). Candida albicans—Biology, molecular characterization, pathogenicity, and advances in diagnosis and control – An update. Microbial Pathogenesis, 117, 128–138. https://doi.org/10.1016/j.micpath.2018.02.028

De Rosa FG, Garazzino S, Pasero D, Di Perri G, Ranieri VM. Invasive candidiasis and candidemia: new guidelines. Minerva Anestesiol. 2009;75:453–45

Haley, L. D. (1961). Yeasts of Medical Importance. American Journal of Clinical Pathology, 36(3), 227–234. https://doi.org/10.1093/ajcp/36.3.227

Dujon, B., Sherman, D., Fischer, G., Durrens, P., Casaregola, S., Lafontaine, I., de Montigny, J., Marck, C., Neuvéglise, C., Talla, E., Goffard, N., Frangeul, L., Aigle, M., Anthouard, V., Babour, A., Barbe, V., Barnay, S., Blanchin, S., Beckerich, J.-M., … Souciet, J.-L. (2004). Genome evolution in yeasts. Nature, 430(6995), 35–44. https://doi.org/10.1038/nature02579

De Rosa Arendrup, M. C., & Perlin, D. S. (2014). Echinocandin resistance: An emerging clinical problem? Current Opinion in Infectious Diseases, 27(6), 484–492. https://doi.org/10.1097/QCO.0000000000000111

Pfaller, M. A., Moet, G. J., Messer, S. A., Jones, R. N., & Castanheira, M. (2011). Geographic variations in species distribution and echinocandin and azole antifungal resistance rates among Candida bloodstream infection isolates: Report from the SENTRY Antimicrobial Surveillance Program (2008 to 2009). Journal of Clinical Microbiology, 49(1), 396–399. https://doi.org/10.1128/JCM.01398-10

Ramage, G., Rajendran, R., Sherry, L., & Williams, C. (2012). Fungal Biofilm Resistance. International Journal of Microbiology, 2012, 1–14. https://doi.org/10.1155/2012/528521

Kuhn, D. M., & Ghannoum, M. A. (2004). Candida biofilms: Antifungal resistance and emerging therapeutic options. Current Opinion in Investigational Drugs (London, England: 2000), 5(2), 186-197.

Murat, P., & Balasubramanian, S. (2014). Existence and consequences of G-quadruplex structures in DNA. Current Opinion in Genetics & Development, 25, 22–29. https://doi.org/10.1016/j.gde.2013.10.012

Varshney, D., Spiegel, J., Zyner, K., Tannahill, D., & Balasubramanian, S. (2020). The regulation and functions of DNA and RNA G-quadruplexes. Nature Reviews Molecular Cell Biology, 21(8), 459–474. https://doi.org/10.1038/s41580-020-0236-x

Endoh, T., Kawasaki, Y., & Sugimoto, N. (2013). Suppression of gene expression by G-quadruplexes in open reading frames depends on G-quadruplex stability. Angewandte Chemie (International Ed. in English), 52(21), 5522–5526. https://doi.org/10.1002/anie.201300058

Balasubramanian, S., Hurley, L. H., & Neidle, S. (2011). Targeting G-quadruplexes in gene promoters: A novel anticancer strategy? Nature Reviews. Drug Discovery, 10(4), 261–275. https://doi.org/10.1038/nrd3428

Warner, E. F., Bohálová, N., Brázda, V., Waller, Z. A. E., & Bidula, S. (2021). Analysis of putative quadruplex-forming sequences in fungal genomes: Novel antifungal targets? Microbial Genomics, 7(5). https://doi.org/10.1099/mgen.0.000570

Meier-Menches, S. M., Neuditschko, B., Zappe, K., Schaier, M., Gerner, M. C., Schmetterer, K. G., Del Favero, G., Bonsignore, R., Cichna-Markl, M., Koellensperger, G., Casini, A., & Gerner, C. (2020). Cover Feature: An Organometallic Gold(I) Bis-N-Heterocyclic Carbene Complex with Multimodal Activity in Ovarian Cancer Cells (Chem. Eur. J. 67/2020). Chemistry – A European Journal, 26(67), 15340–15340. https://doi.org/10.1002/chem.202004459

Morris, G. M., Huey, R., Lindstrom, W., Sanner, M. F., Belew, R. K., Goodsell, D. S., & Olson, A. J. (2009). AutoDock4 and AutoDockTools4: Automated docking with selective receptor flexibility. Journal of Computational Chemistry, 30(16), 2785–2791. https://doi.org/10.1002/jcc.21256

Pires, D. E. V., Blundell, T. L., & Ascher, D. B. (2015). pkCSM: Predicting Small-Molecule Pharmacokinetic and Toxicity Properties Using Graph-Based Signatures. Journal of Medicinal Chemistry, 58(9), 4066–4072. https://doi.org/10.1021/acs.jmedchem.5b00104

Guan, L., Yang, H., Cai, Y., Sun, L., Di, P., Li, W., Liu, G., & Tang, Y. (2019). ADMET-score - a comprehensive scoring function for evaluation of chemical drug-likeness. MedChemComm, 10(1), 148–157. https://doi.org/10.1039/C8MD00472B

