## Supplementary materials_candida_manuscript for "Identification and Targeting Putative G-Quadruplex Sequences in *Candida glabrata*: A Route to Virulence and Pathogenesis Control?" (from another version of this preprint: version 1, 2021-08-08)

Supplementary Figures and Tables:

**Figure S1:** Structure of Compound A, B, and C after optimization

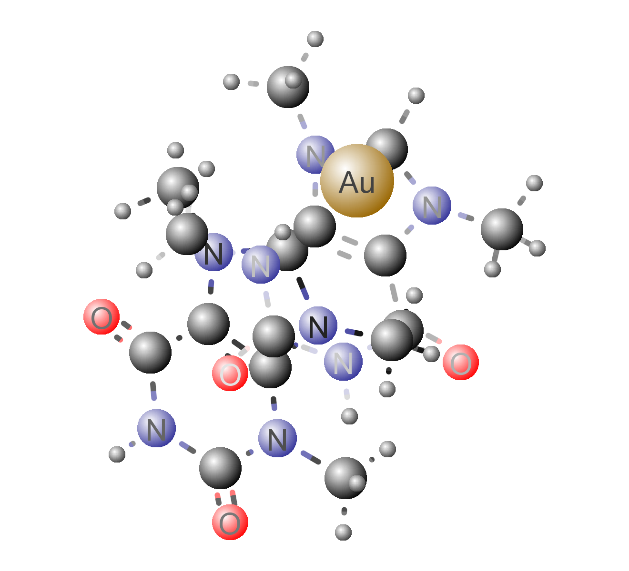

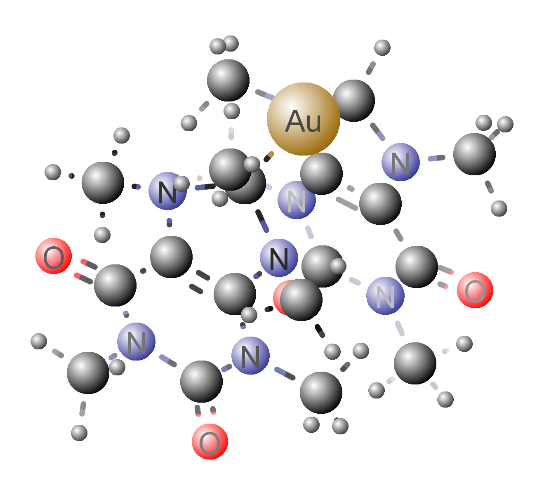

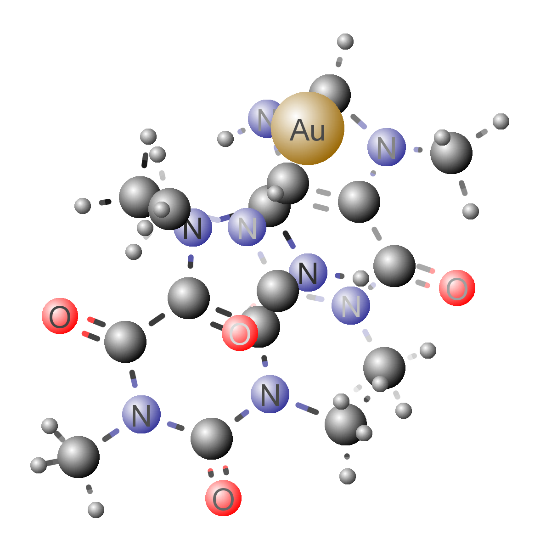

C)

B)

A)

**ST 1.1** depicted the details of the G-quadruplex forming sequence occurring with more than 98% frequency and present in the crucial genes of chromosome A of the strains of Bg2 Candida.

| Systematic Name | Sequence | Probable Molecular Function | Gene Direction | QGRS Score |
| --- | --- | --- | --- | --- |
| CAGL0A04543g | GGGGGGACGGGGAGACGGG | Putative component of the U5 snRNP | Anti-sense Strand | 38 |
| CAGL0A04037g | GGGAAAGGGGAGGGTGGTGGG | Orthologs have role RNA processing | Sense Strand | 40 |
| CAGL0A01243g | GGGGGAGGGGGGCGTGGGGGGG | Predicted Transmembrane transporter activity | Sense Strand | 39 |

**ST1.2** depicted the details of the G-quadruplex forming sequence occurring with more than 98% frequency and present in the crucial genes of chromosome B of the strains of Bg2 Candida.

| Systematic Name | Sequence | Probable Molecular Function | Gene Direction | QGRS Score |
| --- | --- | --- | --- | --- |
| CAGL0B04939g | GGGAGAAGGGAGAAGGGGGATTAGGG | Ortholog to YLR173W | Anti-Sense Strand | 41 |

**ST 1.3** depicted the details of the G-quadruplex forming sequence occurring with more than 98% frequency and present in the crucial genes of chromosome C of the strains of Bg2 Candida.

| Systematic Name | Sequence | Probable Molecular Function | Gene Direction | QGRS Score |
| --- | --- | --- | --- | --- |
| CAGL0C03762g, CAGL0C03740g | GGGGAGGGAACTGGAAGAGGGGGG | Unknown Function | Anti-Sense Strand | 32 |
| CAGL0C05005g | GGGGGAAGGGGGAGGGGGG | Ortholog(s) have protein serine/threonine kinase activity and role in meiotic spindle disassembly, mitotic cytokinesis | Sense Strand | 39 |
| CAGL0C00693g | GGGAGCGGGAGCGGGAGCGGG | Putative Vacuolar Transporter | Sense Strand | 42 |

**ST 1.4** depicted the details of the G-quadruplex forming sequence occurring with more than 98% frequency and present in the crucial genes of chromosome D of the strains of Bg2 Candida.

| Systematic Name | Sequence | Probable Molecular Function | Gene Direction | QGRS Score |
| --- | --- | --- | --- | --- |
| CAGL0D00286g | GGGAAGTGGGGAGGGAAAGGG | Beta mannosyl  transferase activity | Anti-Sense Strand | 40 |
| CAGL0D00286g | GGGGTGGAGGGTGGGGGGAGG | Beta mannosyl  transferase activity | Anti-Sense Strand | 38 |
| CAGL0D00682g | GGGGGGGGCAATGGGGGGAGGGGGG | Predicted nucleic acid binding protein | Anti-Sense Strand | 59 |
| CAGL0D03872g | GGAAGTGGGGGGGGAGGGGGGGGGGG | Unknown Function | Anti-Sense Strand | 62 |
| CAGL0D04400g | GGGGATTAGGACTTGGGTGGGGGG | Orthologs have enzyme inhibitor activity, role in negative regulation of DNA recombinase mediator complex assembly | Anti-Sense Strand | 32 |
| CAGL0D04686g | GGGGGGAAGGGGAAGGGG | Ortholog(s) have metalloendopeptidase activity and role in axial cellular bud site selection | Anti-Sense Strand | 39 |
| CAGL0D05456g | GGGAAGGGGGGGGGGAAGG | Unknown Function | Anti-Sense Strand | 32 |

| Systematic Name | Sequence | Probable Molecular Function | Gene Direction | QGRS Score |
| --- | --- | --- | --- | --- |
| CAGL0D05456g | GGGGGAGGGGGGGGGGGGGGGG | Unknown Function | Anti-Sense Strand | 83 |
| CAGL0D06424g | GGGACGGGGGACGGGGAGGG | Putative aconitase hydratase | Anti-Sense Strand | 41 |
| CAGL0D06512g | GGGGGGGGGGGGGGGGG | Putative membrane bound guanine nucleotide exchange factor | Anti-Sense Strand | 42 |
| CAGL0D04268g | GGGGAGGGGGGGAAGTGGG | Orthologs have role in DNA-dependent DNA replication maintenance of fidelity and Ctf18 RFC-like complex localization | Sense Strand | 39 |
| CAGL0D02244g | GGGAGTGGGGGGACTGGG | Ortholog(s) have protein serine/threonine kinase activity | Sense Strand | 39 |
| CAGL0D00286G | GGGGGAAGAGGGCAGGGAGGGG | Beta Mannosyl transferase activity | Sense Strand | 40 |

**ST 1.5** depicted the details of the G-quadruplex forming sequence occurring with more than 98% frequency and present in the crucial genes of chromosome E of the strains of Bg2 Candida.

| Systematic Name | Sequence | Probable Molecular Function | Gene Direction | QGRS Score |
| --- | --- | --- | --- | --- |
| CAGL0E06182g | GGGAGGGGGAGGGGGGG | Orthologs have role in mitotic sister chromatid segregation and NMS complex | Sense Strand | 41 |
| CAGL0E00671g | GGGTGGGGGGACGGGG | Predicted acid-amino acid ligase activity | Sense Strand | 40 |
| CAGL0E06116g | GGGGAGGGGGGAAGGGAGGG | Orthologs have role in positive regulation of transcription by RNA polymerase II and cytosol | Anti-Sense Strand | 40 |

**ST 1.6** depicted the details of the G-quadruplex forming sequence occurring with more than 98% frequency and present in the crucial genes of chromosome F of the strains of Bg2 Candida.

| Systematic Name | Sequence | Probable Molecular Function | Gene Direction | QGRS Score |
| --- | --- | --- | --- | --- |
| CAGL0F08745g | GGGCTGTGGGGGGGGGGGGAAGGGG | Orthologs have role in cellular response to desiccation and cytoplasm localization | Sense Strand | 42 |
| CAGL0F00605g | GGGGGGAGGGGGGGG | Aldohexose specific glucokinase activity | Sense Strand | 41 |

**ST 1.7** depicted the details of the G-quadruplex forming sequence occurring with more than 98% frequency and present in the crucial genes of chromosome G of the strains of Bg2 Candida.

| Systematic Name | Sequence | Probable Molecular Function | Gene Direction | QGRS Score |
| --- | --- | --- | --- | --- |
| CAGL0G00176g | GGGGGGGGGGGGGG | COPII receptor activity | Anti-Sense Strand | 41 |
| CAGL0G02409g | GGGGGGGGGGGGGGGGG | Orthologs have nucleolus localization | Anti-Sense Strand | 42 |

| Systematic Name | Sequence | Probable Molecular Function | Gene Direction | QGRS Score |
| --- | --- | --- | --- | --- |
| CAGL0G02827g | GGGGATGGGGGATTGGGGATGGG | Ortholog(s) have phosphatidylinositol-4,5-bisphosphate binding, sphingolipid binding activity | Anti-Sense Strand | 39 |
| CAGL0G04455g | GGGAATGGGAGGGGGTGGGGGGG | Ortholog have protein kinase activity, role in negative regulation of DNA replication origin binding | Anti-Sense Strand | 42 |
| CAGL0G08470g | GGGGTGGGGGGATGGGGTGGTGG | Ortholog have DNA topoisomerase activity, DNA topoisomerase type I (single strand cut, ATP-independent) activity | Anti-Sense Strand | 40 |
| CAGL0G08558g | GGGGAACGGGAAAGGGGGGAGG | Orthologs have GTP binding, GTPase activity | Anti-Sense Strand | 39 |
| CAGL0G09064g | GGGGGGGAGGGGGG | Orthologs have role in glycerol biosynthetic process and cytosol, nucleus localization | Sense Strand | 41 |

| CAGL0G05588G | GGGAGTGGGTTGGGTGGG | Ortholog have single-stranded telomeric DNA binding, telomerase inhibitor activity | Sense Strand | 40 |
| --- | --- | --- | --- | --- |
| CAGL0G04719g | GGGGGAAGAGGGGGGGGGGGG | Ortholog have phosphatidylinositol-3,5-bisphosphate 5-phosphatase activity, phosphatidylinositol-3-phosphatase activity | Sense Strand | 39 |

**ST 1.8** depicted the details of the G-quadruplex forming sequence occurring with more than 98% frequency and present in the crucial genes of chromosome H of the strains of Bg2 Candida.

| Systematic Name | Sequence | Probable Molecular Function | Gene Direction | QGRS Score |
| --- | --- | --- | --- | --- |
| CAGL0H08239g | GGGGGGGAAGGGGAGGGAATGG | Ortholog(s) have NAD-dependent histone deacetylase activity | Sense Strand | 41 |
| CAGL0H04895g | GGGGTGGGGGGGGGGG | Ortholog(s) have role in DNA recombinase assembly, error-free translesion synthesis | Sense Strand | 42 |
| CAGL0H02343G | GGGGGGGGGGGGGGGGGGGGGG | Ortholog(s) have transcription corepressor activity | Sense Strand | 83 |

**ST 1.9** depicted the details of the G-quadruplex forming sequence occurring with more than 98% frequency and present in the crucial genes of chromosome I of the strains of Bg2 Candida.

| Systematic Name | Sequence | Probable Molecular Function | Gene Direction | QGRS Score |
| --- | --- | --- | --- | --- |
| CAGL0I03960g | GGGAGGAAGGGGGGGGGGGG | Putative site-specific DNA endonuclease with autocatalytic protein-splicing activity | Anti-Sense Strand | 38 |
| CAGL0I04004g | GGGCGTGGGTGTGGGGGCGGGGGGG | Unknown Function | Anti-Sense Strand | 42 |
| CAGL0I04774g | GGGGGAGGCGGGGGGAAGGGGG | Ortholog(s) have enzyme regulator activity, role in cellular calcium ion homeostasis, glycosphingolipid biosynthetic process | Sense Strand | 38 |
| CAGL0I04994G | GGGGGGTAAGGGGGGG | 5-methyltetrahydropteroyltriglutamate homocysteine methyltransferase | Anti-Sense Strand | 39 |
| CAGL0I05852g | GGAGGGGGGGATGGGAGGTGGGGGGG | Ortholog(s) have cyclin-dependent protein serine/threonine kinase regulator activity | Anti-Sense Strand | 39 |
| CAGL0I07227g | GGGATGGGGGGAAGGG | Putative isocitrate dehydrogenase | Anti-Sense Strand | 40 |
| CAGL0I07821g | GGGGGAGGGGGGATGGCCCTGGGG | Ortholog(s) have ubiquitin binding activity | Anti-Sense Strand | 34 |
| CAGL0I07821g | GGGCTTGGGCTTGGGCTTGGGG | Ortholog(s) have ubiquitin binding activity | Anti-Sense Strand | 42 |
| CAGL0I10147g | GGGGAGGGGGGGGGG | putative adhesin-like protein | Anti-Sense Strand | 41 |
| CAGL0I00506g | GGGGAAGAGGGGGAACGGGATGGG | Ortholog(s) have glutathione hydrolase activity | Sense Strand | 39 |

**ST 1.10** depicted the details of the G-quadruplex forming sequence occurring with more than 98% frequency and present in the crucial genes of chromosome J of the strains of Bg2 Candida.

| Systematic Name | Sequence | Probable Molecular Function | Gene Direction | QGRS Score |
| --- | --- | --- | --- | --- |
| CAGL0J09988g | GGGTGGGCTTGGGCCCGGG | Ortholog(s) have S-adenosylmethionine-dependent methyltransferase activity | Anti-Sense Strand | 40 |
| CAGL0J09702g | GGGGGGGAGGGGGATGGGGGG | Ortholog(s) have role in fungal-type cell wall organization | Sense Strand | 40 |
| CAGL0J09548g | GGGCCTGGGCCTGGGCCTGGG | Ortholog(s) have DNA translocase activity, RNA polymerase II activating transcription factor binding activity | Sense Strand | 42 |

**ST 1.11** depicted the details of the G-quadruplex forming sequence occurring with more than 98% frequency and present in the crucial genes of chromosome K of the strains of Bg2 Candida.

| Systematic Name | Sequence | Probable Molecular Function | Gene Direction | QGRS Score |
| --- | --- | --- | --- | --- |
| CAGL0K12012g | GGGTATGGGTTTGGGCTGTGGG | Lysyl-tRNA synthetase | Ant-Sense Strand | 41 |
| CAGL0K10780g | GGGTGGGAAGAGGGATTGGG | Ortholog(s) have IMP dehydrogenase activity and cytoplasm localization | Anti-Sense Strand | 39 |
| CAGL0K09746g | GGGAGGGAGGGGGTGGGGGGG | Ortholog(s) have role in DNA repair, regulation of transcription by RNA polymerase II and NuA4 histone acetyltransferase complex localization | Sense Strand | 41 |

**ST 1.12** depicted the details of the G-quadruplex forming sequence occurring with more than 98% frequency and present in the crucial genes of chromosome L of the strains of Bg2 Candida.

| Systematic Name | Sequence | Probable Molecular Function | Gene Direction | QGRS Score |
| --- | --- | --- | --- | --- |
| CAGL0L10934g | GGGCAATGGGCGATGGGCGATGGG | Ortholog(s) have role in DNA repair, regulation of transcription by RNA polymerase II and NuA4 histone acetyltransferase complex localization | Ant-Sense Strand | 42 |
| CAGL0L12056g | GGGTTGGGATTTGGGATATGGG | 14-3-3 protein | Anti-Sense Strand | 42 |

**ST 1.13** depicted the details of the G-quadruplex forming sequence occurring with more than 98% frequency and present in the crucial genes of chromosome M of the strains of Bg2 Candida.

| Systematic Name | Sequence | Probable Molecular Function | Gene Direction | QGRS Score |
| --- | --- | --- | --- | --- |
| CAGL0M104174g | GGGCAATGGGCGATGGGCGATGGG | Ortholog(s) have role in DNA repair, regulation of transcription by RNA polymerase II. | Ant-Sense Strand | 41 |
| CAGL0M11748g | GGGGAGGGGGAGGGTAAGGGG | Ortholog(s) have MAP kinase activity, calmodulin binding | Anti-Sense Strand | 41 |
